# Development of microsatellite markers for the threatened species *Coleocephalocereus purpureus* (Cactaceae) using next-generation sequencing

**DOI:** 10.1101/838870

**Authors:** Daphne Amaral Fraga, Anderson Figueiredo de Carvalho, Ricardo Souza Santana, Marlon Câmara Machado, Gustavo Augusto Lacorte

## Abstract

Ten microsatellite loci were developed and validated for the endangered cactus species *Coleocephalocereus purpureus*. The markers were obtained from sequences generated by whole genome shotgun sequencing approaches. A testing group of 36 specimens of the main grouping were genotyped and all described markers presented suitable outcomes to population genetic studies, showing polymorphic status for *C. purpureus* testing group with clean and reproducible amplification. No evidence for scoring errors, null alleles or linkage disequilibrium was detected. Number of alleles per locus ranged from 3 to 6 and expected heterozygosity ranged from 0.78 to 0.99. These new microsatellite loci are suitable to be used in future diversity and structure population studies of *C. purpureus*.

## Introduction

*Coleocephalocereus purpureus* (Buining & Brederoo) F. Ritter is a rare columnar cactus species, reaching up to 90 cm high, with grey wool and golden yellow to brown bristles, that forms a lateral cephalium up to 50 cm long near the stem’s tip with pink melocactus-like flowers (Figure 1) [1]. As a typical columnar cactus, *C. purpureus* specimens provide nectar, pollen and fruits to a broad spectrum of bat, bird, and insect species, playing a crucial support ecological role of a fragile arid ecosystem [2].

**Figure 1.**
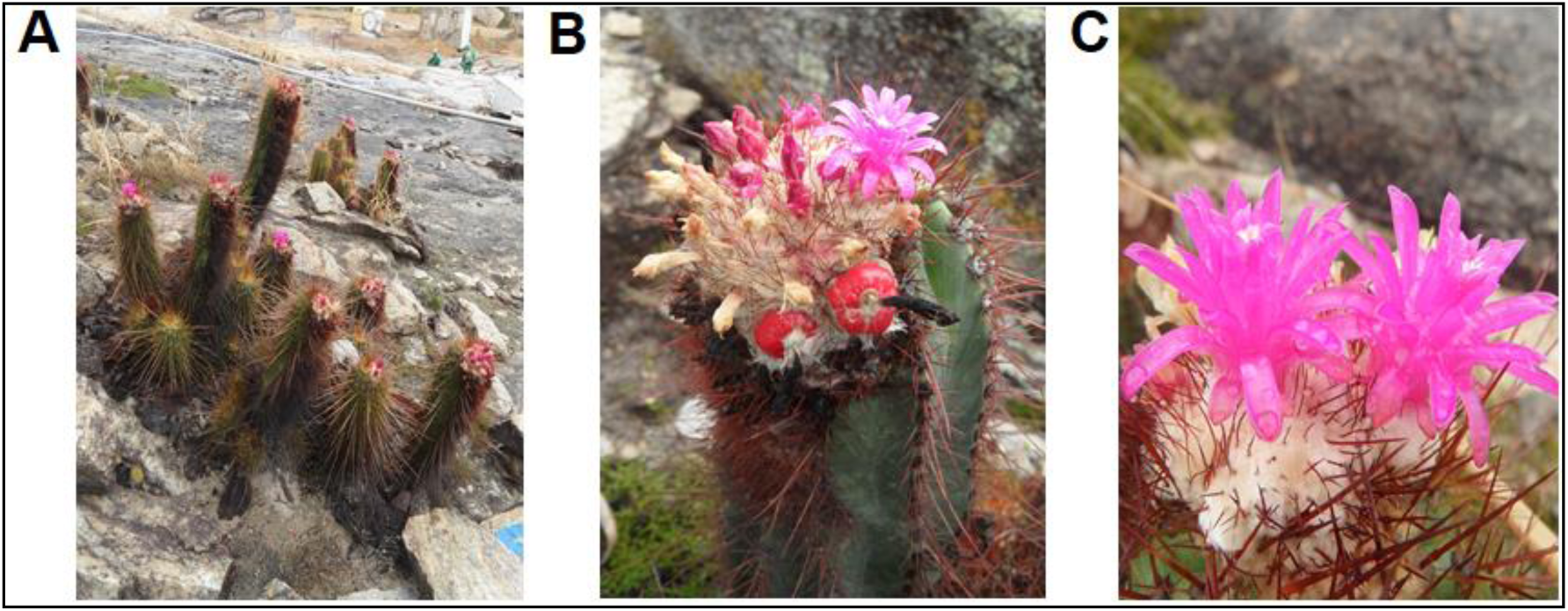
Images of *Coleocephalocereus purpureus.* A, a typical individual of *C. purpureus* with several columnar stems. B, a typical cephalium near the stem’s tip. C, pink melocactus-like flowers.

The species was ranked by IUCN as “critically endangered” due its high endemism, whose occurrence of populations (or even of a single population, the population boundaries is unknown) is limited to only four small habitat patches restricted to inland cliffs and mountain peaks placed in a small area of Caatinga biome from Southeast Brazil (Figure 2), as well due the presence of quarrying activities of the rock where the species grows and other minor human impacts such as collection, deforestation, introduction of invasive species, fire and cattle trampling [3].

**Figure 2.**
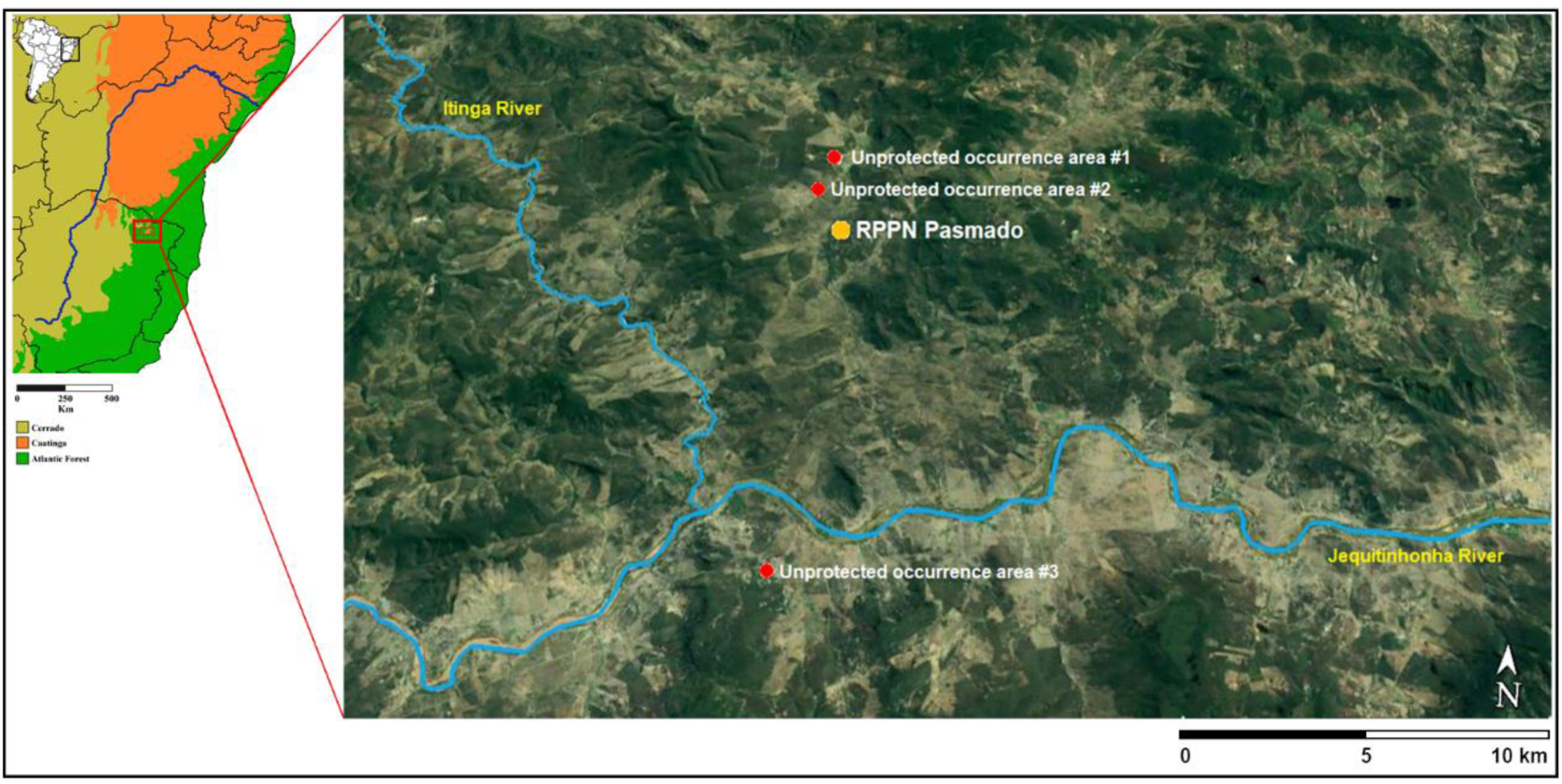
*C. purpureus* distribution. Natural color Google Earth image showing the location of the known occurrence areas of *C. purpureus* remmant individuals. Unprotected areas is represented by red dots and the protected and managed area by a yellow dot. Jequitinhonha and Itinga rivers are depicted in blue. Map of eastern Brazil showing the area studied within the red square (on the left).

Due to the high risk of extinction of the species imposed by quarrying activities in its home range, a conservation unit (named “RPPN Pasmado”) has recently been established to protect one of the main native populations of *C. purpureus* as well as to harbour rescued individuals from areas where quarrying activities has been authorized and all vegetation has been removed [4]. Since the translocation programs were implemented, the RPPN Pasmado Conservation Unit became the largest grouping of *C. purpureus* constituted by native individuals as well translocated individuals from other populational groups that became extinct due to quarrying activity. In addition, as the other three habitat patches harbouring *C. purpureus* individuals are located in unprotected areas of high value for quarrying activities, the RPPN Pasmado Conservation Unit has become the unique expectance of preservation of this species.

Strategies for genetic variability retention of the remaining populations is essential for the persistence of endangered species since low levels of genetic variation decrease population resilience and are associated to fitness decreasing by inbreeding depression [5]. Some genetic studies on cacti species using microsatellite molecular markers have provided insights into genetic diversity, demographic history, population structure and connectivity, which could offer critical guidance for conservation and management strategies [6–8].

The traditional methods for microsatellite isolation were expensive and time-consuming task, drawbacks that often reduced the use microsatellite markers in genetic studies of natural populations [9]. Recently, Next-generation sequencing (NGS), specifically whole genome shotgun sequencing approaches, has become a viable alternative for microsatellite markers isolation since NGS methods comprise faster and less costly approaches besides to generating thousands of markers per procedure [10]. Here, we use whole genome shotgun sequencing approach to develop a set of microsatellite markers for the endemic and endangered species *Coleocephalocereus purpureus* for application in population genetic studies, as a way to guide conservation and management strategies of *C. purpureus* species.

## Material and methods

One *C. purpureus* specimen collected from the RPPN Pasmado was selected for whole genome shotgun sequencing. Total genomic DNA was extracted from a fresh stem slice using NucleoSpin® Plant II DNA extraction kit (Macherey-Nagel, Düren, Germany) and quantified using a Qubit 2.0 device (Life Technologies, Carlsbad, CA, USA). Genomic libraries was constructed using Illumina TruSeq kit (Illumina Inc., San Diego, CA) and were run in a Illumina NextSeq Sequencer (Illumina Inc., San Diego, CA) from BPI Sequencing Facility company (Botucatu, SP. Brazil). The microsatellite motifs were identified from sequences obtained by NGS with MSATCOMMANDER software [11], indicating to the program to search for di, tri, tetra and pentanucleotide microsatellites motifs. Sequencing reads including microsatellite motifs with following specifications were selected to primer design using PRIMER3 [12]: (1) perfect motif microsatellite ≥ 5 tandems repeats; (2) melting temperature around 54°C and with a maximum of 2°C difference between melting temperature for each paired primers; (3) PCR product between 100 and 300 pb length; (4) GC content > 40%; (5) no complementarity between the primers.

A testing group of 36 *C. purpureus* specimens from RPPN Pasmado Conservation Unit were selected and a small stem slice from each individual were collected to DNA extraction using NucleoSpin® Plant II DNA extraction kit (Macherey-Nagel, Düren, Germany). Based on the five defined criteria for primer design, Forty-four primer pair were recovered from PRIMER3 results and subsequently manufactured with 6-FAM™ fluorophore labelling in order to perform PCR optimization and polymorphism tests. The amplification of testing group DNA was performed using a Mastercycler gradient thermocycler (Eppendorf, Germany) and a 25 μL volume reaction containing 50 ng/μL of DNA, 5 μL of PCR Buffer (10 mM Tris–HCl, 50 mM KCl, 1.5 mM MgCl_2_), 0.4 μM of each primer; 0.2 mM dNTPs, 2 units of Taq DNA Polymerase and 6 μL of MiliQ H_2_O. The amplification program included an initial denaturation at 95 °C for 4 min, 30 cycles at 95°C (45 s)/54°C (45 s)/72°C (45 s) and final extension at 72°C for 7 min. Amplification products were visualized in ABI 3730 Genetic Analyzer (Applied Biossystem®) using GeneScan™ 500 LIZ® Size Standard v2.0 (Applied Biossystem®). The genotypes were obtained with Peak Scanner v1.0 software (Applied Biosystems®).

The genetic diversity was evaluated estimating the number of alleles per locus (N_A_), number effective of alleles per locus (N_E_), expected and observed heterozygosity (H_E_ and H_O_, respectively) using the program ARLEQUIN V 3.5.1.2 [13]. In addition, deviations from Hardy–Weinberg equilibrium (HWE) and linkage disequilibrium were tested also using ARLEQUIN V 3.5.1.2 [13]. Frequencies of null alleles were estimated using FREENA [14]. DNA sequences of microsatellite loci described here were deposited at GenBank Database with accession numbers MN200436 to MN200445.

Specimen collection for this study was authorized by Brazilian Environmental Agency (ICMBio) licence number 58257-1. The access to *C. purpureus* genetic material performed in this study was properly registered in official database of Brazilian genetic patrimony – SISGEN – with access number A8AFD21.

## Results and discussion

This is the first time that microsatellite markers for *C. purpureus* were isolated and the NGS approach used in this study generated thousands of sequence reads containing microsatellite motifs (Figure 3). In total, it was generated 17,895,688 reads, of which 14769 sequences containing microsatellite motifs were identified. However, most of sequences containing microsatellite motifs were not suitable to be considered a potential microsatellite molecular marker because they did not present perfect (or even almost perfect) microsatellite motifs, or more than 5 tandems repeats, or sequence length > 100 bp. So, only about two thousand of sequence reads really could be considered potentially amplifiable microsatellite loci.

**Figure 3:**
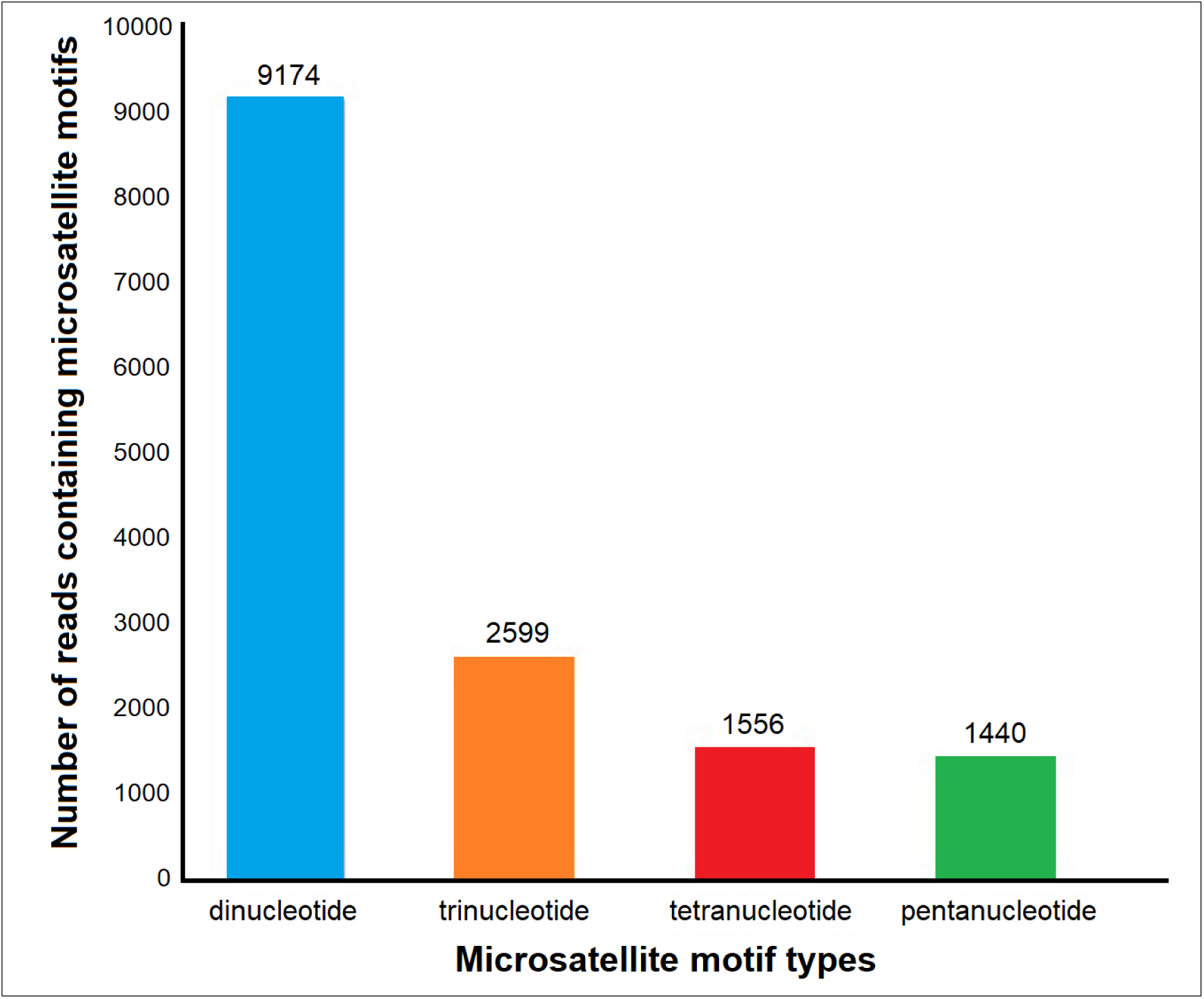
Number of sequence reads containing potentially microsatellite loci (on the top of each bar is the exactly number of reads).

Of all the almost 2,000 potential candidate loci, we selected only those that were accepted by the parameters implemented in the PRIMER3 primer design program, resulting in a list of 44 microsatellite loci. It is important to highlight that in this study we included melting temperature parameter in PRIMER3 software run just to simplify the process of PCR optimization. Thus, more microsatellite loci could have been characterized if the annealing temperature parameter were relaxed or not pre-established. Our results of recovering potentially amplifiable loci, using the whole genome shotgun technique, showed results similar to those performed with the same technique and purpose but for different plant species [15–18]. Our results support the robustness and applicability of NGS approaches for microsatellite loci characterization.

Of the forty-four microsatellite loci selected for the initial tests, ten of them presented suitable outcomes to population genetic studies, showing polymorphic status for *C. purpureus* testing group with clean and reproducible amplification (Table 1). The microsatellite loci presented different patterns of genetic diversity on testing group of 36 *C. purpureus* specimens designed for this study, which the number of alleles per locus ranged from 3 (Cpur-1) to 6 (Cpur-10), the observed heterozysity (Ho) among loci ranged from 0.22 (Cpur-8) to 0.92 (Cpur-3) and expected heterozygosity (H_E_) ranged from 0.78 (Cpur-8) to 0.99 (Cpur-3). Six loci presented HWE deviation attributed to deficit of heterozygotes, which is expected to small populations [19]. No evidence of scoring errors were detected in the genotyping by stuttering or due to large allele dropout as well no evidence of null alleles was verified. No significant linkage disequilibrium was detected.

**Table 1.**
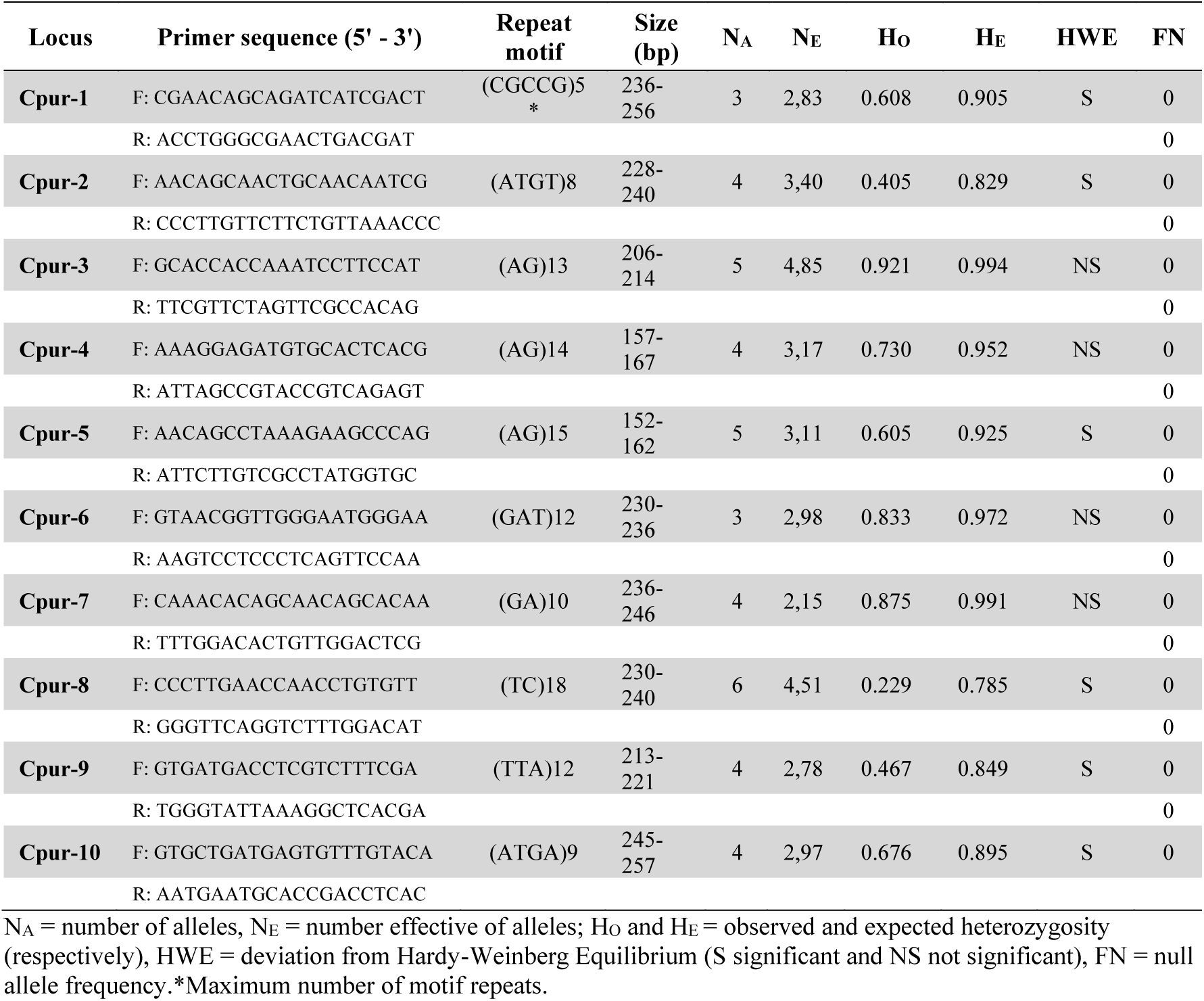
Summary data for 10 microsatellite loci developed for *C. purpureus*

These markers information could be useful for describing the genetic diversity and population structure of the remnant *C. purpureus* populations. They will also be used for understanding patterns of gene flow and potential connectivity among the populations. All this information is crucial for the design of management plans for this threatened species aiming the retention of the residual genetic diversity present in the remnant populations, maximizing the resilience and preventing the effects of inbreeding depression.

## Acknowledgements

The authors thank the Program for Technological Development in Tools for Health-PDTIS FIOCRUZ for use of its facilities and Nativa Serviços Ambientais Ltda for logistic support during collections of biological material.

